# Human Ageing Genomic Resources: 2018 Update

**DOI:** 10.1101/193326

**Authors:** Robi Tacutu, Daniel Thornton, Emily Johnson, Arie Budovsky, Diogo Barardo, Thomas Craig, Gilad Lehmann, Dmitri Toren, Vadim E. Fraifeld, João Pedro de Magalhães

**Affiliations:** Integrative Genomics of Ageing Group, Institute of Ageing and Chronic Disease, University of Liverpool, Liverpool L7 8TX, UK; Computational Biology of Aging Group, Institute of Biochemistry, Romanian Academy, Bucharest, 060031, Romania; The Shraga Segal Department of Microbiology, Immunology and Genetics, Center for Multidisciplinary Research on Aging, Ben-Gurion University of the Negev, Beer-Sheva 84105, Israel; Judea Regional Research & Development Center, Carmel 90404, Israel; Department of Biochemistry, Yong Loo Lin School of Medicine, National University of Singapore, Singapore; Science Division, Yale-NUS College, Singapore

**Keywords:** evolution, genetics, lifespan, life-history

## Abstract

In spite of a growing body of research and data, human ageing remains a poorly understood process. To facilitate studies of ageing, over 10 years ago we developed the Human Ageing Genomic Resources (HAGR), which are now the leading online resource for biogerontologists. In this update, we present HAGR’s main functionalities, including new additions and improvements to HAGR. HAGR consists of five databases: 1) the GenAge database of ageing-related genes, in turn composed of a dataset of >300 human ageing-related genes and a dataset with >2000 genes associated with ageing or longevity in model organisms; 2) the AnAge database of animal ageing and longevity, featuring >4000 species; 3) the GenDR database with >200 genes associated with the life-extending effects of dietary restriction; 4) the LongevityMap database of human genetic association studies of longevity with >500 entries; 5) the DrugAge database with >400 ageing or longevity-associated drugs or compounds; 6) the CellAge database with >200 genes associated with cell senescence. All our databases are manually curated by experts to ensure a high quality data and presented in an intuitive and clear interface that includes cross-links across our databases and to external resources. HAGR is freely available online (http://genomics.senescence.info/).

## Introduction

Ageing is a complex biological process that, despite decades of research, is not yet well understood. Many age-related changes have been described, however the theories regarding what mechanisms drive the changes are still controversial (1). Since their conception, the Human Ageing Genomic Resources (HAGR) have aimed to tackle this complex problem, becoming the leading online resource for biogerontologists. With the advent of large scale sequencing and breakthroughs in the genetics of ageing, HAGR has a particular (but not exclusive) focus on genomics.

As the field of ageing research has grown the amount of data being generated has rapidly increased due to the emergence of high-throughput technologies. Since its first publication in 2005 (2), HAGR has expanded dramatically to match this increase. HAGR organises large quantities of complex data, putting the findings into context and aiding further analysis. Having started with only two databases, GenAge, a database of genes potentially associated with human ageing, and AnAge, a database of ageing, longevity and life history traits in animals (2), HAGR now consists of six databases and a wide range of tools and resources. The different HAGR databases each aim to tackle different aspects of ageing, such as cell senescence, drug and genetic interventions, and genes associated with dietary restriction.

This article provides a non-technical description of the various databases, tools and projects in HAGR and their research applications. New resources created since the 2013 publication (3) are described alongside updates to the remaining resources. In doing so we hope to provide a comprehensive guide to HAGR so they can remain the most accessible and in-depth resources available online in the field of biogerontology. HAGR is freely available online (with no registration required) at http://genomics.senescence.info/.

## Database Content

### GenAge – The Ageing Gene Database

The GenAge database (http://genomics.senescence.info/genes/) is the benchmark database of genes related to ageing. Since its first publication in 2005 (2), GenAge has progressed considerably (Table 1). At first, GenAge only included human genes potentially associated with ageing. Now the database is divided into two main sections: human potential ageing-associated genes and longevity-associated genes in model organisms. In addition, there is also a complimentary list of genes with expression commonly altered during ageing. When the first HAGR paper was published in 2005 (2), GenAge contained 220 entries for human genes. Presently, build 19 (24/06/2017) of GenAge contains 307 human gene entries, 2152 entries for model organisms, and 73 genes with altered expression during ageing. The GenAge databases have previously been described in depth (3), so their use will be only briefly mentioned.

**Table 1.** Growth over time of the various datasets in the GenAge database.

| Database | Species | Number of Gene Entries |  |  |  |
| --- | --- | --- | --- | --- | --- |
|  |  | 2005 | 2009 | 2013 | 2017 |
| GenAge - Human Genes | *Homo sapiens* | 220 | 261 | 288 | 307 |
| GenAge - Model Organisms | *Mus musculus* | - | 68 | 91 | 136 |
|  | *Caenorhabditis elegans* | - | 555 | 680 | 877 |
|  | *Drosophila melanogaster* | - | 75 | 128 | 193 |
|  | *Saccharomyces cerevisiae* | - | 87 | 809 | 909 |
|  | **Total for model organisms** | - | 785 | 1708 | 2115 |

GenAge – Human Genes (http://genomics.senescence.info/genes/human.html) contains a selection of human genes which might affect the ageing process. The focus is on genes implicated in multiple processes and pathologies related to ageing, so those genes affecting only a single age-related process are excluded. Each gene in the dataset is annotated to indicate the strength of the link to human ageing (Table 2). The strongest level of evidence is for those genes directly linked to human ageing, typically those resulting in progeroid syndromes when mutated. So far only three genes (*WRN, LMNA, and ERCC8*) meet these criteria. These data are continuously curated so new results are regularly added to existing entries. Most recent GenAge – Human Genes updates have focused on improving the quality of the data. In addition to new gene entries, older gene entries have been updated to reflect additional findings from new publications.

**Table 2.** Evidence levels linking human genes to ageing, listed from the highest level of evidence to the lowest. Genes with evidence from multiple categories are listed under their highest evidence level.

| Strength of Evidence Linking Human Gene to Ageing | Number of Genes |
| --- | --- |
| Entry selected based on evidence directly linking the gene product to ageing in humans | 3 |
| Entry selected based on evidence directly linking the gene product to ageing in a mammalian model organism | 60 |
| Entry selected based on evidence directly linking the gene product to ageing in a non-mammalian animal model | 25 |
| Entry selected based on evidence directly linking the gene product to ageing in a cellular model system | 16 |
| Entry selected based on evidence linking the gene or its product to human longevity and/or multiple age-related phenotypes | 4 |
| Entry selected based on evidence linking the gene product to the regulation or control of genes previously linked to ageing | 32 |
| Entry selected based on evidence linking the gene product to a pathway or mechanism linked to ageing | 68 |
| Entry selected based on indirect or inconclusive evidence linking the gene product to ageing or showing the gene product to be an effector of genes related to ageing | 99 |

GenAge – Human Genes also includes a list of genes with expression commonly altered during ageing. This list was derived from a meta-analysis of ageing microarray studies in multiple tissues of mammalian species (4). Users can sort the data by Entrez ID, HGNC and gene name. The list is annotated to indicate which genes are overexpressed and which genes are underexpressed.

Using data from GenAge – Human Genes we tracked patterns of ageing research over time. Research into specific genes in the context of ageing mostly began in the 1990s. Certain genes have become well known through their role in ageing. Examples of these include *WRN*, the mutation of which results in Werner syndrome, possibly the most dramatic progeroid syndrome (5), *LMNA*, the mutation of which leads to Hutchinson-Gilford’s progeroid syndrome (6), and *SIRT1*, linked to several processes involved in ageing (7) (Figure 1). For well-studied genes, like *TP53*, an additional role in the ageing process has emerged over time. Examples of these include *MYC*, an oncogene mainly studied in the context of cancer (8), *MTOR*, a regulator of several cellular processes which was found to play a role in ageing in various model organisms (9), and *TP53*, a well-known tumour suppressor (10) (Figure 1).

**Figure 1.**
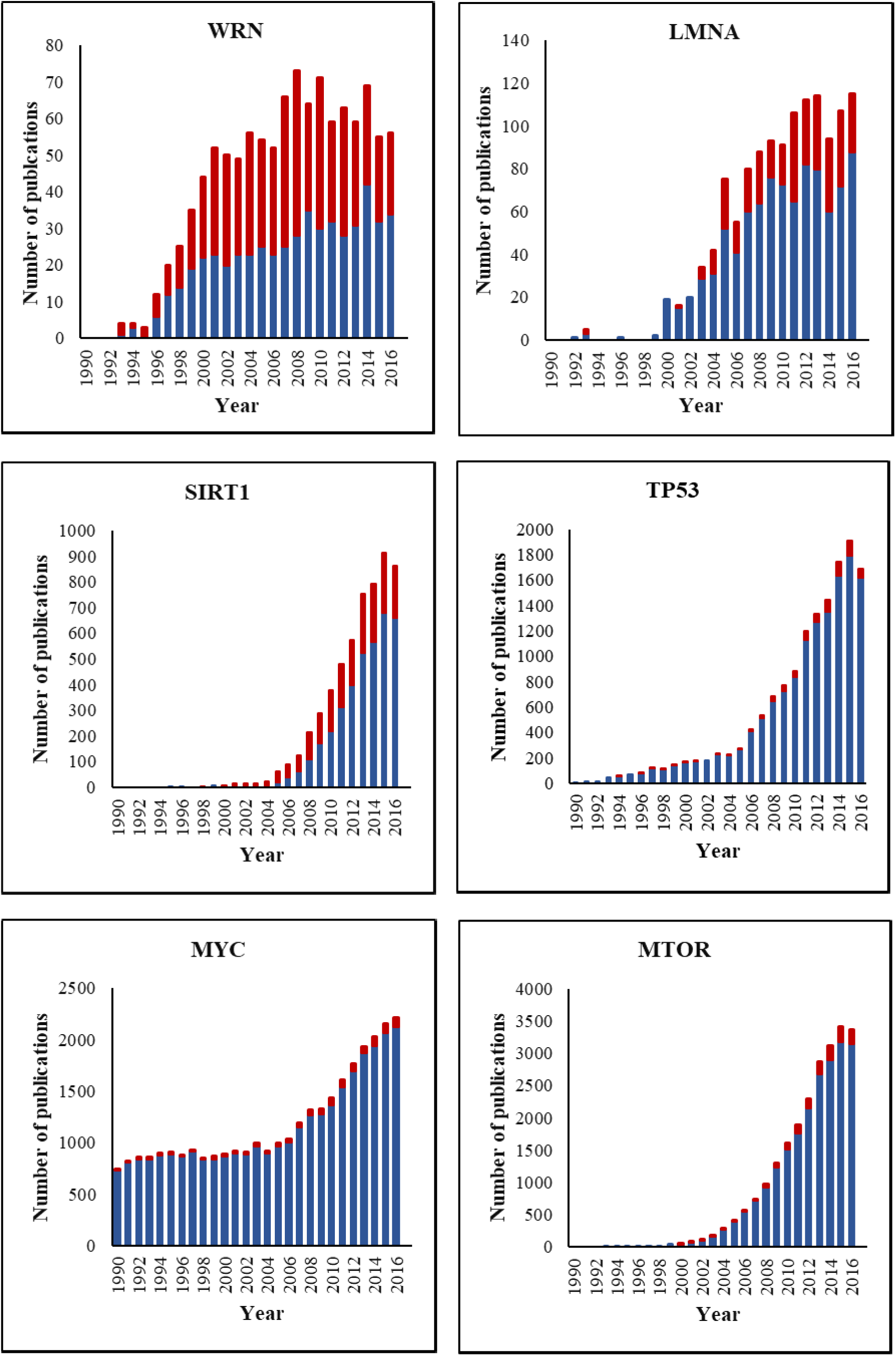
Patterns of ageing research for different human genes over time. The proportion of publications that focused mainly on ageing research are shown in red, the remaining publications are shown in blue.

GenAge – Model Organisms (http://genomics.senescence.info/genes/models.html) is a database of genes, which, if genetically modulated, affect longevity in model organisms (4). Selection was based on gene manipulation experiments (gene knockout, gene knockdown, partial or full loss-of-function mutations and over-expression) which resulted in significant changes in the ageing phenotype and/or lifespan. For studies in mammals a reduction in lifespan was only included if the genetic intervention was specifically linked to ageing. Most observations are from the four most popular biomedical model organisms: mice, worms, fruit flies and yeast (Table 1). However, several results from other model organisms such as zebra fish and golden hamsters are also included. Where reported, the effects of the genetic manipulation on mean and/or maximal lifespan are included to provide quantitative data. As previously detailed (3), genes are categorised as either pro- or anti- longevity, except in large-scale yeast lifespan screens where this can be problematic. Instead pro-longevity genes altering lifespan in large-scale knockout/knockdown yeast lifespan screens are annotated as ‘necessary for fitness’ when a link to ageing has not been made. Where studies report conflicting results, the genes are annotated as ‘unclear’. The size of the model organisms’ dataset has increased almost 3-fold since its conception in 2009.

Users can browse the model organism database using a quick search function. The entire gene list can be sorted by gene symbol, Entrez ID, gene name, organism or maximum changes to the average lifespan. The results can be filtered by species, longevity influence, lifespan effect and key words. For experiments using transgenic organisms, entries are classified according to the species in which the experiments were conducted in, not the species source of DNA. The mouse, worm, fruit fly and yeast data sets are available for download on the model organisms’ homepage. In addition, users can download a list of homologues from other model organisms for each dataset. A list of human homologues for all the genes from model organisms’ genes is also available.

GenAge has proven a valuable resource for ageing research. A systems level analysis of the GenAge human genes database identified a robust group of ageing-specific network characteristics, revealing ageing genes as network hubs and essential genes (11). Moreover, in an analysis of genes in the ageing human brain, 54 genes with sustained, consistent expression and 23 genes with DNA methylation changes were found in GenAge (12). GenAge was also used to validate the targets of a serum miRNA profile of human longevity (13). The data from GenAge has been incorporated into other databases, including the TiRe database (http://www.tiredb.org) (14), a database of tissue repair genes, and the NetAge database (http://netage-project.org) (15), a compendium of networks for longevity, age-related diseases and associated processes.

### AnAge – The Database of Animal Ageing and Longevity

AnAge (http://genomics.senescence.info/species/) is an integrative database of lifespan records for over 4000 organisms. It includes, if available, maximum lifespan, taxonomy, metabolic characteristics, development schedules and a multitude of additional life history data. AnAge was the second database developed for HAGR (2) and is now over a decade old, becoming the most widely used resource in HAGR (see below). It is arguably the ‘gold standard’ for longevity data in animals. Build 14 (15/09/2017) contains 4245 entries.

AnAge has previously been described in depth (3,16), and so its utility will only be briefly described. Users can browse AnAge via the quick search function or manually search all the organisms in the database. Organisms are classified according to Kingdom, Phylum, Class, Order, Family, Genus, and Species. A list of species with negligible senescence is also available. Longevity data contain maximum longevity and, where available, mortality parameters. Entries indicate whether the maximum longevity value comes from specimens kept in captivity or the wild. Each entry includes a qualifier of confidence in the data and an estimate of sample size (3,16). Anecdotal evidence is not used to estimate maximum longevity but may be included in the observations. Factors that might introduce bias into comparative ageing studies such as body size, metabolic rate, and development schedules are also included where available (3,4,16).

Recent updates for AnAge have mostly been qualitative. The rate of new species added has reduced over time but older entries are kept up to date with new findings in the field. Build 14 included ∼150 new references. Whilst the main focus of AnAge remains on data in animals, particularly chordates, the database contains entries for several reference plants and fungi. Entries for fungi are currently limited to three commonly studied model organisms: baker’s yeast, fission yeast and filamentous fungus.

The primary goal of AnAge is as a data source for comparative and evolutionary biogerontological studies, thus enabling researchers to study what factors influence differences in phenotype and longevity across phylogeny. Previously, a comparative study from our group (17) used data from AnAge to show that basal metabolic rate does not correlate with longevity in eutherians and birds and that age at maturity is proportional to lifespan. Data from AnAge has also been used to demonstrate that the largest differences in longevity tends to occur between orders in mammals and genera in birds (18). A recent study demonstrated that brain size and longevity were positively correlated in birds and that major diversification of brain size preceded diversification of longevity in avian evolution. Data on maximum lifespan, body mass, incubation length and clutch size were compiled from AnAge to facilitate this work (19).

Additionally, AnAge has proven a valuable resource for studies in other areas of research. Some examples include determining the relationship between host longevity and parasite species richness in mammals (20) and calculating the effects of radiation dose on wildlife (21). The data from AnAge have also been incorporated into the Comparative Cellular and Molecular Biology of Longevity Database (http://genomics.brocku.ca/ccmbl/) (22), the MitoAge database (http://www.mitoage.info) (23), the Encyclopedia of Life (http://eol.org/), and the Animal Diversity Web (http://animaldiversity.ummz.umich.edu), demonstrating the versatility of this resource.

### GenDR – a Database of Dietary Restriction Related Genes

Dietary restriction (DR) delays the ageing process and extends lifespan in a multitude of species from yeast to mammals such as mice and monkeys (24). However, the exact mechanisms of how DR extends lifespan are still unknown. As previously described (25), GenDR (http://genomics.senescence.info/diet/) is a database of DR-related genes. Herein, the use and function of GenDR will be briefly outlined along with updates since the 2013 paper.

DR-essential genes are defined in GenDR as those, which, if genetically modified, interfere with DR-mediated lifespan extension (3,25). Users can browse gene manipulations in GenDR by organism, with entries for nematodes, fruit flies, mice, budding yeast and fission yeast. For each model organism, a list of DR genes is available for download, along with a list of orthologues in other common model organisms and humans. GenDR also contains a complimentary dataset of genes consistently differentially expressed in mammals under DR (26). Build 4 (24/06/2017) of GenDR contains 214 genes inferred from genetic manipulations and 173 genes from expression changes. GenDR is fully integrated in HAGR with crosslinks to the other databases.

GenDR is the first and only database of dietary restriction genes. We hope that GenDR may aid in the production of pharmacological DR mimetics. Analysis of the gene network of DR genes by our group predicted several novel DR essential genes, which were experimentally validated in yeast (25). GenDR was also used to validate the gene targets of candidate DR mimetics in worms (27). In an analysis of the downstream targets of *daf-16*, a gene involved in DR in worms, four of the targets overlapped with the GenDR database, demonstrating the involvement of different components of the pathway in DR (28).

### LongevityMap – Human Longevity Genetic Variants

Variation in human lifespan has been found to be 20-30% heritable, with increasing heritability at advanced ages (29). To demonstrate the diversity of human genetic variants associated with longevity and catalogue the increasing volume of data, we created LongevityMap (http://genomics.senescence.info/longevity/), a database of genes, gene variants and chromosomal locations associated with longevity. This differs from the GenAge database which focuses mostly on data from model organisms and the few genes associated with human progeroid syndromes. LongevityMap has been described in depth elsewhere (30), so will only be briefly outlined here.

Entries in LongevityMap were compiled from the literature (30). Negative results are included to provide information regarding each gene and variant previously studied in the context of human longevity. Both large and small-scale studies are included, along with several cross-sectional studies or studies of extreme longevity. Due to the diversity of data, details about the study design are outlined for each entry, such as population and sample size (30). Build 3 (24/06/2017) of LongevityMap contains 550 entries, 884 genes and 3144 variants. Of the 550 entries, 275 are reported as significant findings. Users can sort entries by ID, gene/variant, chromosome location, publication date and population. Results can also be filtered to only include certain populations or significant/non-significant findings.

As next generation sequencing and genome-wide approaches advance, so does the capacity for longevity association studies. For example, a functional enrichment study of LongevityMap data yielded clusters, which consisted of regulation of apoptosis, response to environment, response to hormone stimulus, regulation of locomotion and regulation of phosphorylation (31). With similar studies, we hope that LongevityMap will act as a reference to help researchers parse the increasing quantities of data related to the genetics of human longevity.

### DrugAge – a Database of Ageing-related Drugs

Studies performed on model organisms have revealed that ageing is a plastic process that can be manipulated by genetic and environmental factors (32). Thus, identifying drugs that could extend lifespan in model organisms has received considerable interest (33). The DrugAge database (http://genomics.senescence.info/drugs/) is a curated database of drugs, compounds and supplements with anti-ageing effects that extend longevity in model organisms. Build 2 (01/09/2016) of DrugAge was recently described in depth elsewhere (34).

DrugAge was developed to allow researchers to prioritise drugs and compounds relevant to ageing, providing high-quality summary data in model organisms. As described (34), the data were primarily compiled from the literature, in addition to other databases and submissions from the scientific community. Users can browse the database via compound name, dosage, species or lifespan effect. Drug interactions can be browsed by compound name, interaction data source (35,36) and protein identifiers. The current build contains 418 distinct compounds across 1316 lifespan assays on 27 unique model organisms. DrugAge is integrated with other HAGR databases and crosslinks drug-gene interactions to ageing-related genes, allowing users to gain a deeper understanding of the pathways involved in ageing.

Hundreds of genes in several pathways act as regulators of ageing (32). However, analysis of DrugAge and other HAGR databases has revealed that the overlap between the targets of lifespan-extending drugs and known ageing related genes is modest (34). This indicates that most ageing-related pathways have yet to be targeted pharmacologically; DrugAge may aid in guiding further assays. This was recently demonstrated in one study where machine learning was used to predict whether a compound would increase lifespan in worms using data from DrugAge. The best model had 80% prediction accuracy and the top hit compounds could broadly be divided into compounds affecting mitochondria, inflammation, cancer, and gonadotropin-releasing hormone (37). These compounds could all be targets for experimental validation.

### CellAge – a Database of Cell Senescence Genes

Cell senescence, also known as cellular senescence (CS), is the irreversible cessation of cell division of normally proliferating cells. Senescent cells accumulate as an organism ages and may be an important contributor to ageing and age-related disease (38). However, the connection between organismal ageing and CS remains controversial (39,40). CellAge (http://genomics.senescence.info/cells/), a database of CS-associated genes, was created to better understand the mechanisms of CS and its role in ageing.

To create CellAge, a list of CS-associated genes was manually curated from the literature. Selection was based on gene manipulation experiments in human cells, which caused cells to induce or inhibit CS. The type of CS (replicative, stress-induced, oncogene-induced), cell line, cell type and manipulation methods were recorded and can be used to search for records of interest. The database includes data from primary cells in addition to immortalised cell lines and cancer cell lines. Each record contains observations about the evidence. Where reported, common markers of CS (41) such as growth arrest, increased SA-β-galactosidase activity, SA-heterochromatin foci, a decrease in BrdU incorporation, changes in morphology, and specific gene expression signatures are described.

The first build of CellAge contains 279 entries, in which experiments in lung fibroblasts, embryonic kidney cells and foreskin fibroblasts are the most widely represented in the data. It is hoped that CellAge will aid in understanding the various types of CS and that analysis of the data will lead to the discovery of further CS-associated genes. Analysis of the CellAge dataset is currently being carried out by our group (unpublished results). A more detailed description of CellAge and its analysis will be published in a future publication. As for our other databases, CellAge is fully integrated with other HAGR databases and each gene has cross-links to entries in other databases where possible.

## Tools, Projects and Other Information Resources

### Ageing-related Disease Genes

In industrialized societies, ageing is the main risk factor for many debilitating and life-threatening diseases including cancer, cardiovascular disease, arthritis, diabetes, and neurodegeneration. As lifespan increases so too does the prevalence of these diseases (42). An understanding of how these diseases are linked to the ageing process is needed to help tackle this growing problem. The ageing-related disease genes tool (http://genomics.senescence.info/ageing_disease/) makes available a set of age-related disease genes from a 2016 paper (31).

The genes were assembled using data compiled by a National Institute of Ageing study (43), which is available online (https://www.irp.nia.nih.gov/branches/rrb/dna/gene_sets.htm/). Diseases with fewer than 20 genes associated were excluded from the gene list to avoid the dilution of findings. Processes and conditions such as insulin resistance and hyperlipidaemia were classified as dysfunctions and excluded from the list. Users can browse genes and diseases by MeSH disease terms, MeSH disease class and by gene symbol. The disease classes are cardiovascular diseases, immune system diseases, musculoskeletal diseases, neoplasms, nervous system diseases, and nutritional and metabolic diseases. Results can be grouped by gene or disease. There are 769 genes associated with 20 age-related diseases in total.

Our tool was designed so that age-related disease genes can be viewed, analysed and downloaded in the context of ageing genes to understand potential functional overlap. Besides, our tool allows users to create a merged data set between age-related disease genes and ageing genes, according to user-defined filters. Users first select an age-related disease gene set, then select an ageing gene set from one or more of the HAGR databases. The genes are then merged to retrieve common ageing and age-related disease genes. Where applicable, HAGR genes can be converted into human homologues before merging (Figure 2).

**Figure 2.**
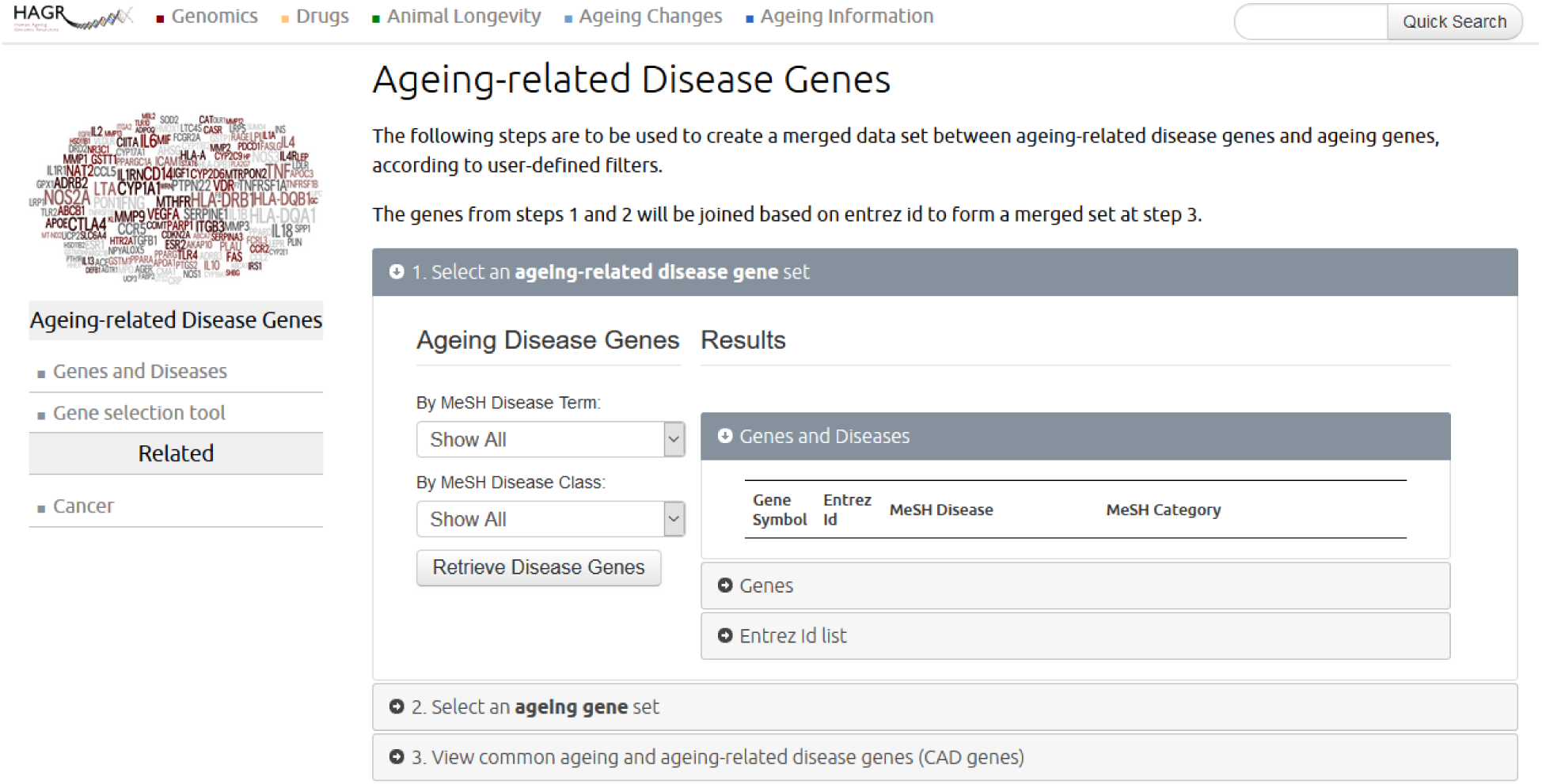
Snapshot of tool for gene analysis of age-related disease genes and their overlap with ageing-related genes.

### Software and Other Datasets

Within HAGR there are several additional projects and resources. During the development of HAGR a series of software programs were created to aid in a variety of bioinformatics analyses. The Ageing Research Computational Tools (ARCT) are a toolkit of Perl modules aimed at parsing files, data-mining, and searching and downloading data from the Internet (2,4). An SPSS script is also available, which can be used to determine the demographic rate of ageing for a given population. This method has been detailed and applied in previous work (44). These resources are available on the HAGR website (http://genomics.senescence.info/software/).

HAGR also makes available multiple datasets relevant to the evolution of ageing (http://genomics.senescence.info/evolution/) and cancer incidence across organs (http://genomics.senescence.info/cancer/).

### Complementary resources to HAGR

For a greater understanding of ageing there are several additional resources to HAGR. The Digital Ageing Atlas (DAA, http://ageing-map.org/) is a database of human age-related changes at different biological levels (45). The DAA integrates molecular, pathological, physiological and psychological age-related data to characterise the human ageing phenotype. The DAA is freely available online and has over 3000 age-related changes associated with specific tissues. The DAA features mouse data to compliment the information on human ageing. HAGR links to the DAA on the homepage. Quick searches within HAGR also show results from the DAA where available.

Senescence.info (http://www.senescence.info) is an informational repository on the science of ageing which aims to highlight the importance of ageing research and give an overview of current knowledge on the biology and genetics of ageing. This resource is targeted at both scientists and non-scientists alike and features numerous tutorials aimed at students and newcomers to the field. Unlike HAGR, senescience.info is developed by a single person (J.P.M).

The WhosAge database (http://whoswho.senescence.info/) is a non-exhaustive list of individuals and companies working on ageing and lifespan extension. Presently, WhosAge features 26 companies and 291 researchers and includes a brief description of the work of each company or individual and relevant contact information for them (4).

Lastly, two social media resources have been made available on Facebook (https://www.facebook.com/pg/BiologyAgingNews) and Twitter (https://twitter.com/AgingBiology) which report on the latest news and findings in the field. The updates detail research in longevity, life-extension and rejuvenation technologies and link to articles/papers for further reading. These resources usually post several times a week to >6,000 followers.

## Availability

Our access policy remains the same as in our previous publications (3). All HAGR databases and resources are freely available at http://genomics.senescence.info/. All databases allow users the opportunity to export, download and reuse data for their own analyses, under a Creative Commons Attribution licence. Feedback is welcome and encouraged via email.

## Concluding Remarks

Over the last decade HAGR has expanded to include several new databases, datasets, tools and additional resources. HAGR emphasizes high quality data on ageing and the databases are under continuous curation by experts in the field. Though the selection process for inclusion of genes in HAGR is kept as objective as possible, some subjectivity is unavoidable. To cope with this, our policy is to be inclusive, providing evidence and links to the relevant literature and thus providing a balanced and comprehensive overview to the reader (4). AnAge provides information on data quality and sample size and prioritises the reliability of the data over the most extreme values. GenAge – Model organisms, GenDR and CellAge all focus on genes from genetic manipulation experiments to ensure the selection process is as unbiased as possible. In doing so we hope users may draw their own conclusions from the data.

Out of all the HAGR resources AnAge remains the most popular (Figure 3). GenAge – Human genes and GenAge – Model Organisms have collectively maintained high levels of use. Since DrugAge was released in 2016 its usage has greatly increased, becoming one of the most widely used databases. CellAge is the newest HAGR resource, released in late 2016, hence not surprisingly still one of the least popular.

**Figure 3.**
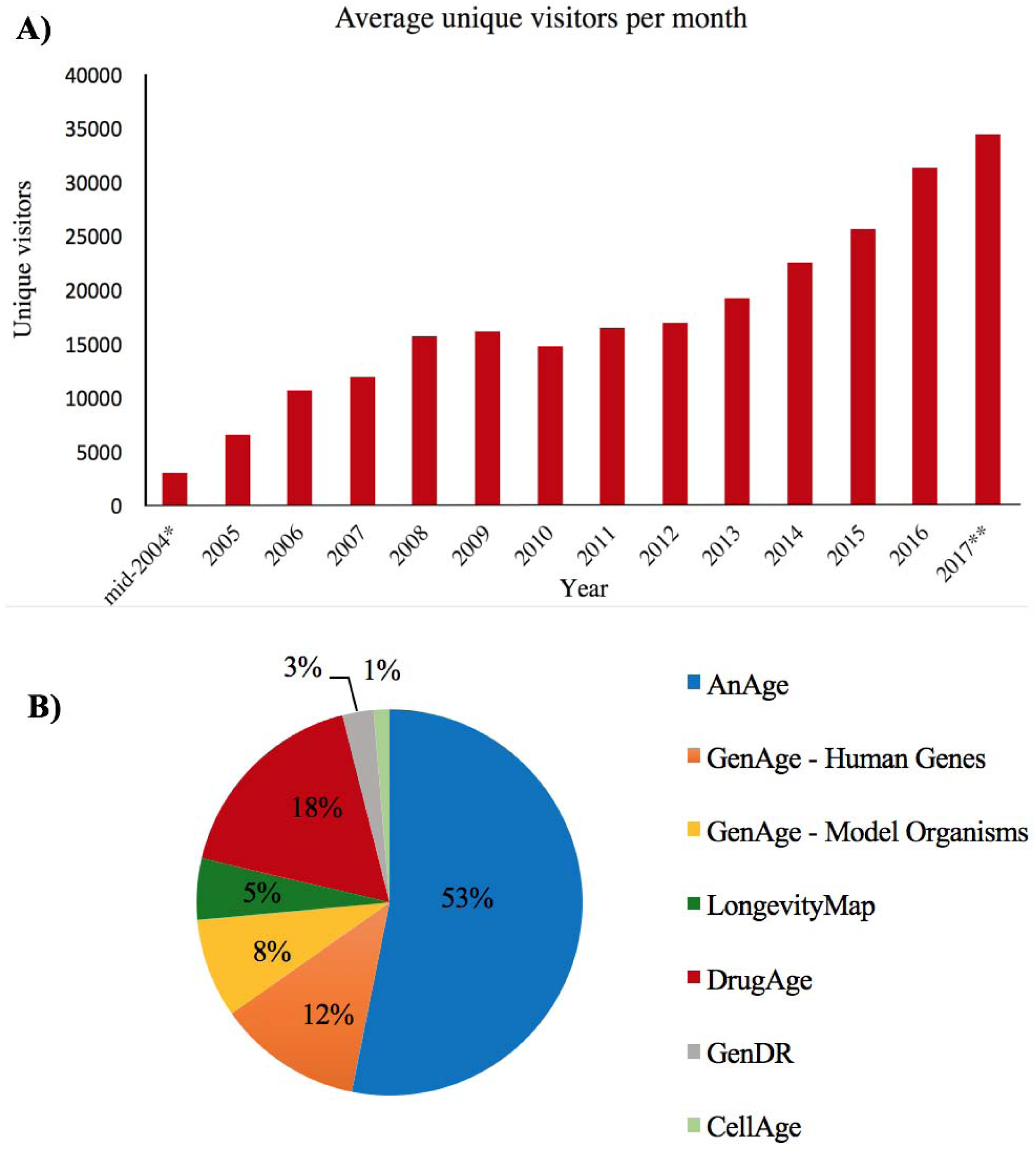
**A) Combined HAGR and** senescence.info unique visitors per month (data to end of August 2017). **B)** Usage by % of the different HAGR databases in 2017.

HAGR has been cited over 500 times since it was first published in 2005 (Table. 3) and has seen a continuous rise in the number of citations over recent years. From 2006 our resources received over,000 unique visitors per month, and they now receive over 30,000 unique visitors per month, thus indicating HAGR’s growing importance in the field.

**Table 3.** Summary of third-party works using/citing HAGR from 2004-2016.

| **Year** | **HAGR citations** |
| --- | --- |
| 2004 | 1 |
| 2005 | 8 |
| 2006 | 7 |
| 2007 | 11 |
| 2008 | 26 |
| 2009 | 33 |
| 2010 | 44 |
| 2011 | 47 |
| 2012 | 43 |
| 2013 | 57 |
| 2014 | 65 |
| 2015 | 85 |
| 2016 | 72 |
| **Total:** | **499** |

HAGR covers many aspects of ageing, acting as a science of ageing portal aimed at an audience from beginners to experts in biogerontology. Visitors are encouraged to send feedback and propose enhancements/features they would like to see in future. Over time as data continues to be generated we anticipate that HAGR will continue to grow to meet this influx, maintaining its status as the leading online resource for biogerontologists.

## Acknowledgements

We would like to thank the many people who have helped in this project, providing valuable advice and suggestions, including former and current members of the Integrative Genomics of Ageing Group. HAGR is supported by a Wellcome Trust grant (104978/Z/14/Z) to JPM. RT is supported by an EU-funded grant, through Competitiveness Operational Programme 2014-2020, POC-A.1-A.1.1.4-E-2015.

